# The “amphi”-brains of amphipods: New insights from the neuroanatomy of *Parhyale hawaiensis* (Dana, 1853)

**DOI:** 10.1101/610295

**Authors:** Christin Wittfoth, Steffen Harzsch, Carsten Wolff, Andy Sombke

**Author notes:** shared last authorship (Carsten Wolff, Andy Sombke).

## Abstract

**Background:** Over the last years, the amphipod crustacean *Parhyale hawaiensis* has developed into an attractive marine animal model for evolutionary developmental studies that offers several advantages over existing experimental organisms. It is easy to rear in laboratory conditions with embryos available year-round and amenable to numerous kinds of embryological and functional genetic manipulations. However, beyond these developmental and genetic analyses, research on the architecture of its nervous system is fragmentary. In order to provide a first neuroanatomical atlas of the brain, we investigated *P. hawaiensis* using immunohistochemical labelings combined with laser-scanning microscopy, X-ray microcomputed tomography, histological sectioning and 3D reconstructions.

**Results:** As in most amphipod crustaceans, the brain is dorsally bent out of the body axis with downward oriented lateral hemispheres of the protocerebrum. It comprises almost all prominent neuropils that are part of the suggested ground pattern of malacostracan crustaceans (except the lobula plate and projection neuron tract neuropil). Beyond a general uniformity of these neuropils, the brain of *P. hawaiensis* is characterized by a modified lamina (first order visual neuropil) and, compared to other Amphipoda, an elaborated central complex. The lamina displays a chambered appearance that, in the light of a recent analysis on photoreceptor projections in *P. hawaiensis*, corresponds to specialized photoreceptor terminals. The presence of a poorly differentiated hemiellipsoid body is indicated and critically discussed.

**Conclusions:** Although amphipod brains show a general uniformity, when compared with each other, there is also a certain degree of variability in architecture and size of different neuropils. In contrast to other amphipods, the brain of *P. hawaiensis* does not display any striking modifications or bias towards one particular sensory modality. Thus, we conclude that its brain may represent a common type of an amphipod brain.

## BACKGROUND

Amphipod crustaceans display a high disparity in body plans, life history, and ecology. Therefore, they are suitable organisms to explore adaptive changes of organ systems, e.g. the nervous system, in response to different life styles. *Parhyale hawaiensis* (Dana, 1853) (Peracarida, Amphipoda, Hyalidae) is an epibenthic amphipod with circumtropical distribution that occupies intertidal marine habitats such as bays, estuaries, and mangroves [1–3] and is also a typical member of the macroalgal fauna [4]. It was first described from the Hawaiian islands [5]. As most representatives of the Hyalidae, these animals show continuous reproduction throughout the year and can adapt their reproduction to favorable environmental conditions [6,7]. Dynamics and demographic parameters of a population in its native range showed two main reproductive periods, a shorter one, from late autumn to early winter, and a longer one, from late spring to early summer [7]. The sex ratio in natural populations of this species typically is biased toward females thus allowing for a rapid increase in abundance when environmental conditions are favorable [8]. Females have a low number of eggs, between six and 25 per brood, depending on the size and age of the females [9]. Because *P. hawaiensis* tolerates salinities from 5 up to 40 PSU [10] and has such a wide distribution, Artal et al. [11] suggested this species as a suitable ecotoxicity test organism for circumtropical nearshore marine ecosystems and described standardized procedures for laboratory husbandry.

In recent years, *P. hawaiensis* has also evolved into an important laboratory model species [12,13]. Its embryogenesis has been thoroughly described including a staging system [14], and fate map and cell lineage analyses of the early embryo [15] until gastrulation [16] were carried out. Because laboratory husbandry is easy and affordable, inbred lab cultures can provide ample material for developmental studies year-round. Furthermore, *P. hawaiensis* is accessible for experimental manipulation and robust protocols exist for the fixation of embryos [17], *in situ* hybridization to study mRNA localization [18], and immunohistochemistry to study protein localization [19]. Genetic tools and resources which have been established in *P. hawaiensis* in recent years include for example stable transgenesis [20–22], gene knockdown [23–25], CRISPR-mediated gene editing [26], transcriptomic approaches [27–29], and a sequenced genome [30]. Individuals (embryonic stages and adults) are also optically tractable providing the opportunity to capture the cellular events contributing to appendage development and regeneration using cutting-edge live-imaging technologies [31,32]. The recent paper by Ramos et al. [33] provided the basis of genetics-driven analysis of visual function in this species.

In comparison to our current understanding of the brain structure in decapod crustaceans [34–37], our knowledge on the nervous system in Peracarida has not kept pace and for the Amphipoda relies on older studies including those by Gräber [38] and Hanström [39]. Exceptions include representatives of the genus *Gammarus* in which the structure of the ventral nerve cord [40] and brain [41] have been explored in detail including immunohistochemical techniques [42]. These previous reports already described that, within Amphipoda, the brain is dorsally bent out of the neuraxis so that the protocerebrum is almost tilted backwards. The brain in representatives of the genus *Orchestia* was analysed by Madsen [43] and, in comparison to the amphipod *Niphargus puteanus*, by Ramm and Scholtz [44], the latter study using a set of contemporary neuroanatomical techniques, which is comparable to that used in the present report. Ramm and Scholtz [44] provided a detailed description of brain neuropils and soma clusters that will serve as a sound basis to which we compare our own results. Gross anatomy of the central nervous system of *P. hawaiensis* was already documented in drawings by Divakaran [45] who unfortunately did not provide any micrographs. Ramos et al. [33] analysed the structure of the compound eyes and retinal projections in *P. hawaiensis*. Our investigation sets out to explore the neuroanatomy of this emerging crustacean model organism in detail with a set of complementary techniques including classical histology, immunohistochemistry and confocal laser-scan microscopy, x-ray microscopy, and three-dimensional reconstruction. Therefore, as first neuroanatomical atlas of the brain of *P. hawaiensis*, the current report aims to provide the basis for subsequent studies to gain deeper insights into the neurobiology of this emerging model organism, such as functional studies and connectomics.

## METHODS

### Experimental animals

Specimens of *Parhyale hawaiensis* (Dana, 1853) were reared in aquaria with artificial seawater (32 PSU) at about 26 °C. For all experiments, pairs in precopula (Fig. 1A) were collected to ensure maturity of both sexes.

**Fig. 1:**
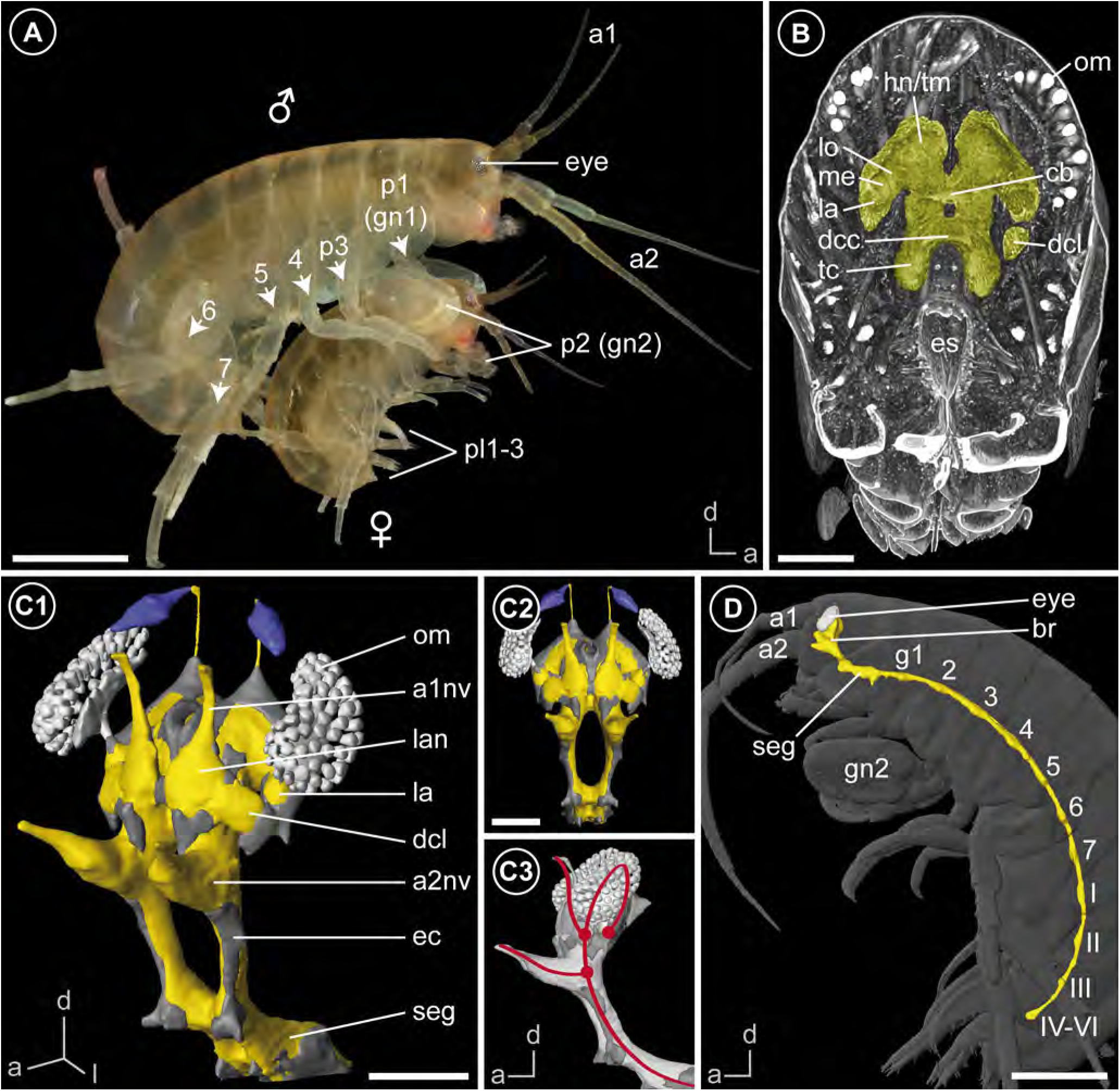
The amphipod *Parhyale hawaiensis* and gross morphology of its nervous system. (**A**) Male and female of *P. hawaiensis* in precopula. (**B**) Orthoslice at the mid-level of the brain based on microCT analysis. The brain (yellow) is situated right between the compound eyes in the dorsal part of the head. (**C**) 3D reconstruction of the brain based on microCT analysis (light grey: optical system, yellow: neuropil, purple: organ of Bellonci, dark grey: somata) in anterolateral view (C1), frontal view (C2) and lateral view (C3). The red line displays the neuraxis of the brain with red dots indicating the roots of associated nerves to sensory organs and appendages to signify the arrangement of proto-, deuto-, and tritocerebrum. An interactive 3D PDF is available as supplement (Additional file 1). (**D**) 3D reconstruction of the central nervous system (yellow) in anatomical context based on microCT analysis in lateral view. The central nervous system is located ventrally and bent dorsally, anterior to the subesophageal ganglion. *<u>Abbreviations:</u> I-III* segmental ganglia of the pleosome, IV-VI fused ganglion of the urosome, *a1* antenna 1, *a1nv* antenna 1 nerve, *a2* antenna 2, *a2nv* antenna 2 nerve, *br* brain, *cb* central body, *dcc* deutocerebral commissure, *dcl* deutocerebral chemosensory lobe, *ec* esophageal connectives, *es* esophagus, *g1-*7 segmental ganglia of the pereon, gn2 second gnathopod, *hn/tm* hemiellipsoid body/ terminal medulla complex, *la* lamina, *lan* lateral antenna 1 neuropil, *lo* lobula, *me* medulla, *om* ommatidia, *p1-7* pereopods 1-7, *pl1-3* pleopods 1-3, *seg* subesophageal ganglion, *tc* tritocerebrum. *<u>Scale bars:</u>* (A, D) 1mm, (B) 200 µm, (C) 150 µm.

### Immunohistochemistry

Three different marker sets were chosen for immunohistochemical labeling. A summary of used protocols is given in table 1 and the reagents used are listed in table 2. Additional notes are given below:

**Tab. 1:** Used protocols for immunohistochemical labelings.

| vibratome sections (100 μm) |  | whole-mounts |
| --- | --- | --- |
| tubulin + FMRFamide | tubulin + histamine | synapsin |
| anaesthetise at 4 °C |  |  |
| 1. FIXATION |  |  |
| Bouin's fixative<br>until usage at 4 °C<br>(at least overnight) | 4 % EDAC in PBS + 1 % DMSO<br>+ 5 % glucose<br>for 30 min. + 24 h at 4 °C | 3 % glyoxal<br>overnight at 4 °C<br>(following Richter et al. 2018) |
|  | post-fixation: 4 % PFA<br>for 24 h at 4 °C |  |
| 2. PREPARATION |  |  |
| decapitation | decapitation | dissect brain |
| VIBRATOME – SECTIONING |  |  |
| wash in PBS and transfer into poly-L-lysine<br>embed in gelatine-ovalbumin<br>post-fix in 1:20 formaldehyde-PBS at 4 °C |  |  |
| 3. PRIMARY ANTISERA |  |  |
| wash in PBTx (PBS + 0.5 % Triton X-100 + 1 % bovine serum albumin)<br>for 2 x 15 min + 60 min at room temperature |  |  |
| 1:1000 anti-acetylated-tubulin<br>1:1000 anti-FMRFamide | 1:1000 anti-acetylated-tubulin<br>1:1000 anti-histamine | 1:1000 anti-SYNORF1 synapsin |
| 1.5 days at room temperature |  | 3.5 days at room temperature |
| 4. SECONDARY ANTISERA |  |  |
| wash in PBTx (0.1 M PBS + 0.5 % Triton X-100 + 1 % BSA)<br>for 2 x 10 min + 2 x 20 min at room temperature |  |  |
| 1:500 Cy3 anti-mouse<br>1:500 Alexa488 anti-rabbit<br>1:1000 Hoechst 33258 |  | 1:500 Cy3 anti-mouse<br>1:1000 Hoechst 33258 |
| 1.5 days at room temperature |  | 2.5 days at room temperature |
| 5. POSTPROCESSING |  |  |
| wash in several changes of 0.1 M PBS (for at least six changes in 3 hours) |  |  |
| mount in Mowiol 4-88 (Roth 0713.2) |  | 1:1 glycerine-PBS for 20 min<br>4:1 glycerine-PBS for 60 min<br>mount in 4:1 Dabco-glycerine |

**Tab. 2:** Used reagents for immunohistochemical labelings

| Labeling reagent | Supplier and specification |
| --- | --- |
| monoclonal anti-acetylated α-tubulin antibody<br>produced in mouse | Sigma T6793, [1] |
| polyclonal anti-FMRFamide antiserum<br>produced in rabbit | Acris/Immunostar 20091, [1,2] |
| polyclonal anti-histamine antiserum<br>produced in rabbit | Progen 16043, [2–4] |
| monoclonal anti-SYNORF1 synapsin antibody<br>produced in mouse | DSHB 3C11, [1,3] |
| polyclonal Cy3 anti-mouse IgG secondary antibody<br>produced in goat | Jackson Immuno Research 115-165-003,<br>[1] |
| polyclonal Alexa 488 anti-rabbit IgG secondary antibody<br>produced in goat | Invitrogen A11008, [1,3] |
| Hoechst 33258 | Sigma 14530, [1] |

- When using 4 % paraformaldehyde (PFA) as fixative, small crystals occurred around the brain, which masked the specific signal during confocal laser-scanning microscopy. In order to avoid this artefact, we used Bouin’s fixative [46] and 3 % glyoxal [47] for immunohistochemical labeling against anti-acetylated-tubulin and anti-SYNORF1 synapsin, respectively.
- Using Bouin’s fixative for immunohistochemical labeling required several changes of 0.1 M PBS (phosphate buffered saline, pH 7.4) to wash out the picric acid completely. A final washing step was carried out overnight.
- The brain of *P. hawaiensis* is embedded within connective tissue rich in lipids (Fig. 2A). Therefore, for immunohistochemical labelings against anti-histamine, we added 1 % DMSO to 4 % EDAC in PBS to increase penetration ability through fatty tissues [48].

### Histology

For section series, three adult individuals (two males, one female) were decapitated and prefixed for 24 h in a solution of ten parts 80 % ethanol, four parts 37 % formaldehyde and one part 100 % acetic acid (compare [49]). After washing in PBS, specimens were postfixed for 1 h in 2 % OsO_4_ solution (same buffer) at room temperature and, following dehydration in a graded series of acetone, embedded in Araldite (Araldite CY212; Agar Scientific #AGR1030). Serial semi-thin sections (1 or 1.5 µm) were prepared with a Microm HM 355 S and stained using 1 % toluidine blue and PyroninG in a solution of 1 % sodium tetraborate.

**Fig. 2:**
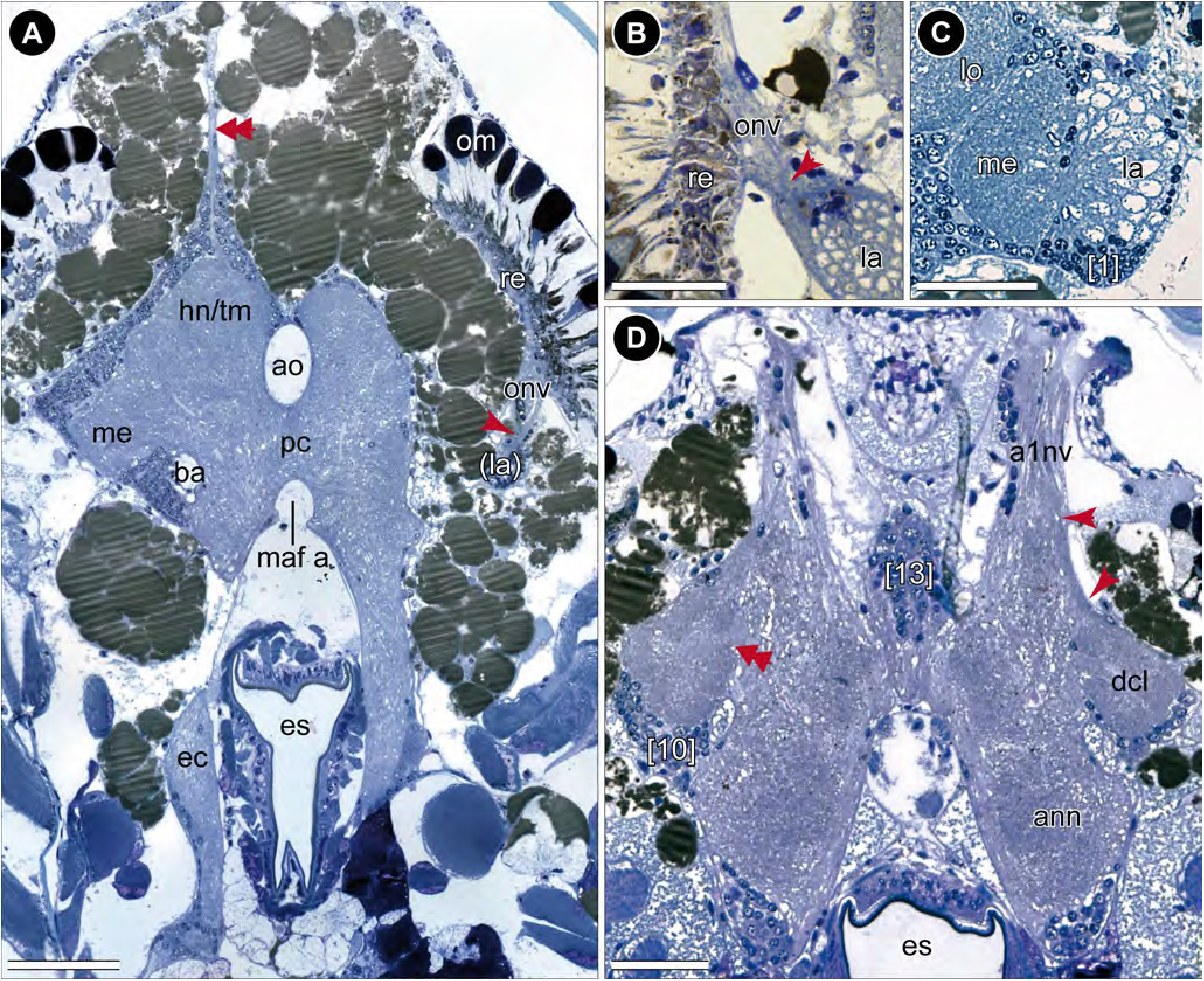
Histological sections of the head. (**A**) Frontal section of the head with posterior part of the brain. The brain is surrounded by numerous lipid droplets (brownish spherules). The dorsal-most part of the brain is innervated by paired small nerves (double-arrow). The optical nerve forms a chiasm (red arrow). (**B**) Magnification of the chiasm (red arrow) of the optical nerve of the left brain hemisphere, frontal view. (**C**) Magnification of the first and second visual neuropil of the right brain hemisphere displaying a chambered appearance of the lamina, frontal view. Numerous terminations within each chamber are visible. Somata of cluster 1 appear darker and smaller. (**D**) Frontal section of the anterior part of the brain. A small section of the antenna 1 nerve projects to the deutocerebral chemosensory lobe (red arrows). The medial foramen is discernable (double-arrow). *<u>Abbreviations:</u> a1nv* antenna 1 nerve, *ann* antenna 2 neuropil, *ao* anterior aorta, *ba* brain artery, *dcl* deutocerebral chemosensory lobe, *ec* esophageal connective, *es* esophagus, *hn/tm* hemiellipsoid body/ terminal medulla complex, *la* lamina, *lo* lobula, *maf a* myoarterial formation a, *me* medulla, *om* ommatidia, *onv* optical nerve, *pc* protocerebrum, *re* retina, *[numbers]* somata cluster. *<u>Scale bars:</u>* (A) 100 µm, (B-D) 50 µm.

### X-ray micro-computed tomography

After cold anesthetization, several specimens were fixed in Bouin’s solution overnight. The subsequent preparation followed the protocol by Sombke et al. [50]. Preparations were rinsed in several changes of PBS, dehydrated in a graded ethanol series and incubated in a 1 % iodine solution (iodine resublimated in 99 % ethanol; Carl Roth #864.1) for 8–12 hours. Preparations were rinsed several times in pure ethanol and critical-point-dried (Leica EM COD300). Finally, samples were fixed on insect pins with super glue. MicroCT scans (n=9) were performed with a Zeiss XRadia MicroXCT-200 and analyzed. One overview scan (young male, 4x objective lens unit, pixel size 5.6521 µm) and two detailed scans of the head (two males, 10x and 20x objective lens units, pixel sizes 1.9649 µm and 0.9983 µm, respectively) were used for visualization and reconstruction in this contribution. Tomography projections were reconstructed using the XMReconstructor software (Zeiss Microscopy) resulting in image stacks (TIFF format). All scans were performed using binning 2 (resulting in noise reduction) and subsequently reconstructed using binning 1 (full resolution) to avoid information loss.

### Imaging

Immunohistochemical preparations were analyzed with a Leica TCS SP5 II confocal laser-scanning microscope equipped with DPSS-, Diode- and Argon-lasers and operated by the Leica Application Suite Advanced Fluorescence software package (LASAF). Images of single frames and maximal projections were compiled using the software LASAF (Leica Microsystems CMS GmbH) and image processing platform Fiji [51].

Specimens processed for histology were analyzed with a Nikon Eclipse 90i upright microscope and bright-field optics (20x objective).

The precopula was photographed with a Canon 70d camera equipped with an EF-S 18-135 mm f/3.5-5.6 IS objective and a Macro Twin Lite MT-24EX flashlight. Cross-polarized light was used to minimize reflections [52].

### Elastic alignment and 3D-reconstruction

Mosaic image data of histological sections were compiled into single image stacks using Adobe Bridge CS4 combined with the Photomerge function of Adobe Photoshop CS4. To perform 3D-reconstructions, an elastic alignment was performed using the plugins Elastic Stack Alignment and Elastic Montage incorporated in the TrakEM2 software of FIJI [53].

3D-reconstructions were prepared with the software Amira 5.4.3 (FEI Visualization Sciences Group, Thermo Fisher Scientific). For using the Segmentation Editor in Amira, the aligned image stacks were converted into greyscale images. Contours of the brain and selected subunits were traced in single section images and finally used to calculate 3D surface models. For 3D-reconstruction of microCT scans, image stacks of virtual sections were processed in the same way. Additionally, based on semi-thin sections, we counted the number of nuclei in the brain cortex of one hemisphere of one adult male and one adult female. For this purpose, Amira’s Filament Editor was used.

### Presentation of data and terminology

Images were processed with Adobe Photoshop CS4 using global picture enhancement features (i.e. brightness and contrast). The diagram was created with Adobe Illustrator CS4. Unless indicated otherwise, all images are oriented with dorsal to the top and, on lateral views, anterior facing to the left. Local arrangement of all neuronal structures, the neuropils and tracts, are described referring to the body axis.

The neuroanatomical nomenclature is based on Sandeman et al. [54] and Richter et al. [55] with modifications adopted from Loesel et al. [56] and Kenning et al. [57] for the description of brain neuropils, cell clusters and tracts. Hence, we name the visual neuropils lamina, medulla, and lobula and use the terms ‘deutocerebral chemosensory lobe’ instead of ‘olfactory lobe’ [57] and ‘projection neuron tract’ instead of ‘olfactory globular tract’ [56]. The arteries are classified after Wirkner and Richter [58].

## RESULTS

### Gross morphology

The head of *Parhyale hawaiensis* is flattened anteriorly. The bilaterally paired and sessile compound eyes are reniform and located dorsolaterally at the head capsule (Fig. 1A, D). Between the compound eyes, the pair of uniramous, short first antennae (a1) are located at the anterodorsal edge of the head capsule. The pair of uniramous second antennae (a2) are nearly twice as long as the first antennae and located ventrally at the mid-level of the head. The complex feeding apparatus consisting of mandibles, first and second maxillae as well as maxillipeds follows ventrally (Fig. 1A, D).

The central nervous system is composed of the brain (br) and ventral nerve cord (vnc), the latter comprises a fused subesophageal ganglion (seg), seven segmental ganglia of the pereon (g 1-7), three segmental ganglia of the pleosome (g I-III), and one fused ganglion of the urosome (g IV-VI, Fig. 1D). The neuraxis of the ventral nerve cord follows the body axis, but the axis of the brain neuromeres is bent anterodorsally (Fig. 1C3). In consequence and corresponding to the anterodorsally situated head appendages, the brain of *P. hawaiensis* is located in the anterodorsal part of the head, between the compound eyes (Fig. 1B-D; Add. file 1). The three neuromeres of the brain, proto-, deuto-, and tritocerebrum, are lined up from dorsal to ventral with the lateral protocerebrum facing posteroventrally towards the ventral-most level of the compound eyes (Fig. 1C3). The neuromeres of the brain are highly fused, but can be distinguished by their associated nerves from sensory organs and appendages. The compound eyes are connected with the protocerebrum *via* the optical nerve (onv, Figs. 2A, 3A). Additionally, the dorsomedial area of the protocerebrum is associated with a bilaterally paired, small nerve that originates in the organ of Bellonci (Fig. 1C2, double arrow in Figs. 2A, 8; Add. file 1). The antenna 1 nerve (a1nv) innervates the deutocerebrum from anterodorsal (Figs. 1C, 2D, 3B, 6B), while the antenna 2 nerve (a2nv) innervates the tritocerebrum from anterior (Figs. 1C, 3B). The shape of the brain is strongly influenced by four large arteries, which proceed anteriorly through the protocerebrum: the (1) median anterior aorta (ao), the (2) median myoarterial formation A (maf a) which branches off anteriorly and, a (3) bilaterally paired, smaller brain artery (ba), located between the lateral and median protocerebrum (Figs. 2A, 3, 8).

### Protocerebrum

#### Lateral protocerebrum

The lateral extensions of the protocerebrum (lateral protocerebrum) are confluent with the median protocerebrum. The anterior aorta separates dorsomedially the hemiganglia of the protocerebrum (Figs. 1B, 2A, 3A, 8). From ventrolateral to dorsomedial, the lateral protocerebrum consists of the serially arranged visual neuropils lamina (la), medulla (me), and lobula (lo) as well as the hemiellipsoid body (hn) and terminal medulla (tm; also termed ‘medulla terminalis’; Figs. 1B, 4A, 7; Add. files 2 and 3). Lamina and medulla are medially separated from the median protocerebrum by the paired brain arteries (ba), whereas lobula, hemiellipsoid body, and terminal medulla are fused ventrally with the anteromedial protocerebral neuropil (ampn, Figs. 3A, 4A). The retina (re) is connected to the first order visual neuropil, the lamina, *via* a short optical nerve (onv), which forms a chiasm (arrow in Fig. 2A, B). In comparison to the size of the compound eyes, the lamina is a relatively small, compact structure. Synapsin-like immunoreactivity reveals a cup-shaped architecture composed of spherical subunits that do not appear to be arranged in an ordered fashion (Fig. 6A, A1). Correspondingly, in anti-acetylated α-tubulin labelings (Fig. 4A) and histological sections (Fig. 2C), the lamina exhibits a chambered structure with a considerable number of small terminations from retinula cell axons within each subunit (Fig. 2C). Comparable to synapsin-like immunoreactivity, a strong histamine-like immunoreactivity is mostly concentrated in the periphery of these spherical subunits (asterisks in Fig. 4C1, C2). Furthermore, anti-histamine labelings reveal that the lamina is organized into two layers with a thin inner (proximal) layer displaying a stronger immunoreactivity than a thick outer (distal) layer (brackets in Fig. 4C2). The second visual neuropil, the medulla, is reniform and anteroventrally enclosed by the lamina (Figs. 2C, 4A, B, C, 6A). Between lamina and medulla, the outer chiasm is formed by a considerable number of neurites (arrow in Fig. 4B; Figs. 2C, 4C1). Adjacent to the medulla and medially fused to the anteromedial protocerebral neuropil, the lobula (the third order visual neuropil) is hardly discriminable. However, the lobula can be discerned by its connection to the central body (cb, see below) *via* a small tract (double-arrows in Figs. 3A, 4A). In sagittal sections, several crossing neurites between medulla and lobula are discernable which are suggested to represent the inner chiasm (arrow in Fig. 4D). In the dorsal-most part of the brain, the hemiellipsoid body and terminal medulla form a complex (Figs. 2A, 3A, B, 4A, 5A, 6A). These two neuropils are not clearly distinguishable by most methodological approaches used. Synapsin-like immunoreactivity reveals a slight distinction of a cap-shaped area in the medial-most part of the brain, which may represent the hemiellipsoid body (Fig. 6A2). However, anti-acetylated α-tubulin labeling reveals slight differences in intensity of immunoreactivity in two equally sized regions (Figs. 4A, 5A) and another approximate distinction is evident by a diffuse signal medially in preparations immunolabeled against RFamide-like peptides (Figs. 3A, 4A). Therefore, the hemiellipsoid body may also be larger than anticipated in anti-SYNORF 1 synapsin labelings.

**Fig. 3:**
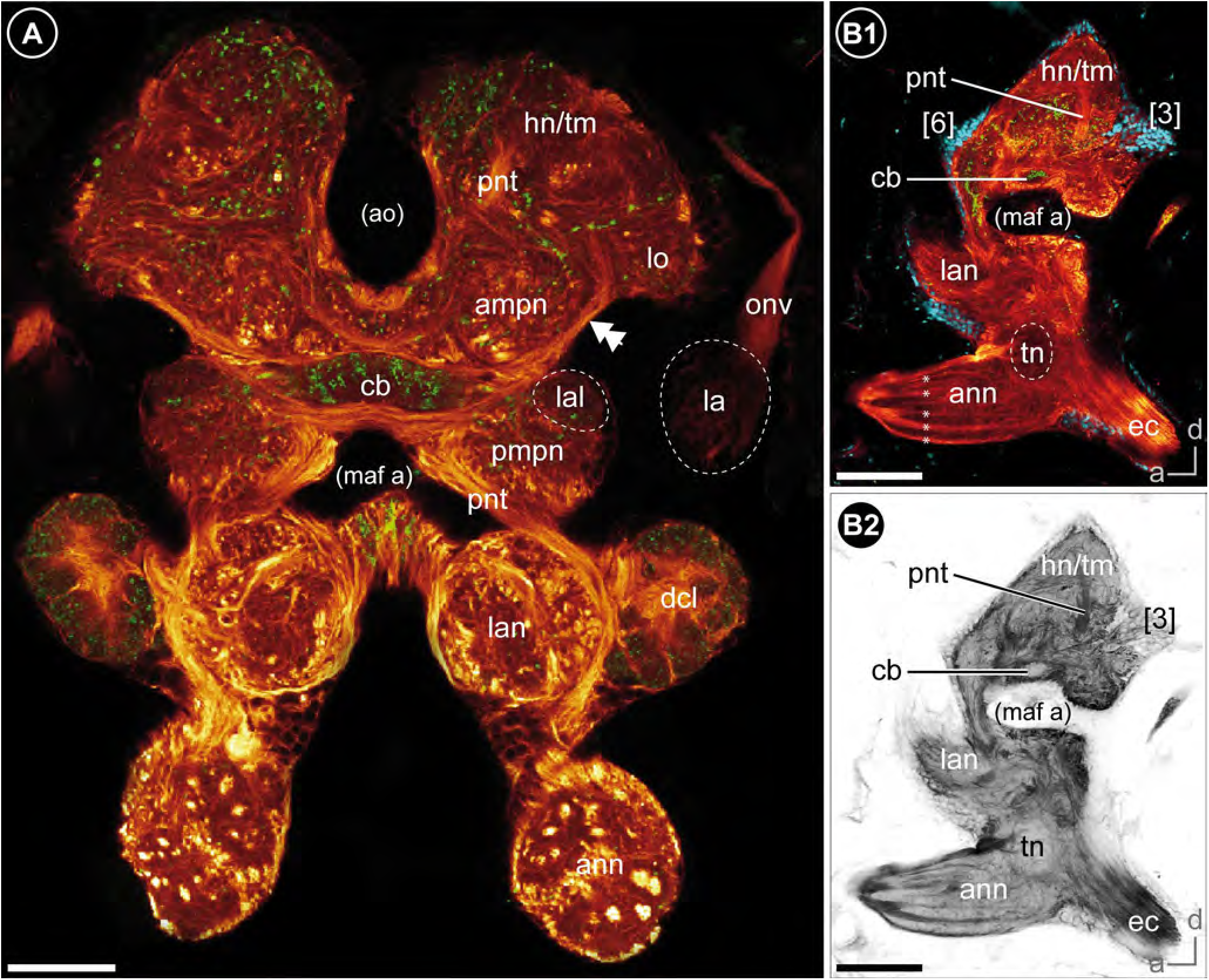
Overview of the neuroanatomy of the brain. (**A**) Frontal vibratome section at the mid-level of the brain, double-labeled against acetylated tubulin (yellow-red) and RFamide (green). (**B**) Sagittal vibratome section of the brain, triple-labeled against acetylated tubulin (yellow-red), RFamide (green) and nuclei (blue, B1) as well as against acetylated tubulin, separately (black-white inverted, B2). *<u>Abbreviations:</u> ampn* anterior medial protocerebral neuropil, *ann* antenna 2 neuropil, *ao* anterior aorta, *cb* central body, *dcl* deutocerebral chemosensory lobe, *ec* esophageal connective, *hn/tm* hemiellipsoid body/ terminal medulla complex, *la* lamina, *lal* lateral accessory lobe, *lan* lateral antenna 1 neuropil, *lo* lobula, *maf a* myoarterial formation a, *onv* optical nerve, *pmpn* posterior medial protocerebral neuropil, *pnt* projection neuron tract, *tn* tegumentary neuropil, *[numbers]* somata cluster. *<u>Scale bars:</u>* (A) 50 µm, (B) 100 µm.

**Fig. 4:**
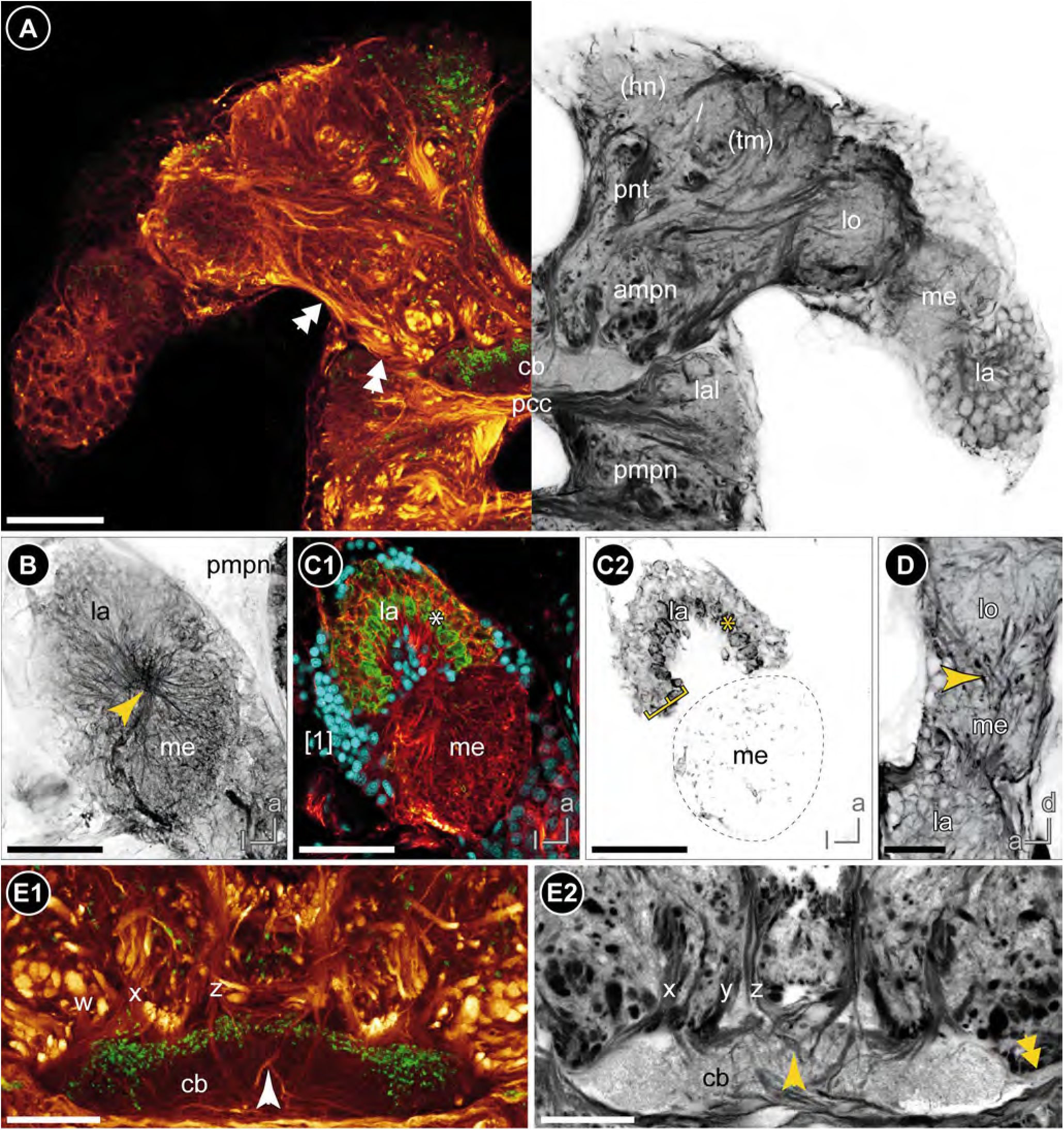
The protocerebrum. (**A**) Frontal vibratome section at the mid-level of the protocerebrum, double-labeled against acetylated tubulin (yellow-red on the left, black-white inverted on the right) and RFamide (green on the left). A bundle of neurites connects the lobula with the lateral part of the central body (double-arrows). (**B-C**) Horizontal vibratome section of the first and second visual neuropils of the left brain hemisphere. (B) Maximal projection of acetylated tubulin-like immunoreactivity (black-white inverted) showing the outer chiasm between lamina and medulla (yellow arrow). (C) Single optical slice, triple-labeled against acetylated tubulin (yellow-red), histamine (green) and nuclei (blue, C1) as well as against histamine, separately (black-white inverted, C2). Histamine-like immunoreactivity is concentrated peripherally within the spherical subunits of the lamina (asterisks) and reveals a two-layered organization of the lamina (brackets in C2). (**D**) Sagittal vibratome section of the visual neuropils showing crossing neurites between lamina and medulla (outer chiasm) as well as between medulla and lobula (putative inner chiasm, yellow arrow), labeled against acetylated tubulin (black-white inverted). (**E**) Frontal view on the central body and the tracts w, x, y, and z, double-labeled against acetylated tubulin (yellow-red), RFamide (green, E1), as well as against acetylated tubulin, separately (black-white inverted, E2). Some neurites of z-tracts form chiasmata within the central body (single arrow). The double-arrow points to the bundle of neurites that connects the central body with the lobula. *<u>Abbreviations:</u> ampn* anterior medial protocerebral neuropil, *cb* central body, *hn/tm* hemiellipsoid body/ terminal medulla complex, *la* lamina, *lal* lateral accessory lobe, *lo* lobula, *me* medulla, *pcc* protocerebral commissure, *pmpn* posterior medial protocerebral neuropil, *pnt* projection neuron tract, *w/x/y/z* tracts, *[numbers]* somata cluster. *<u>Scale</u> <u>bars:</u>* (A-C) 50 µm, (D-E) 25 µm.

**Fig. 5:**
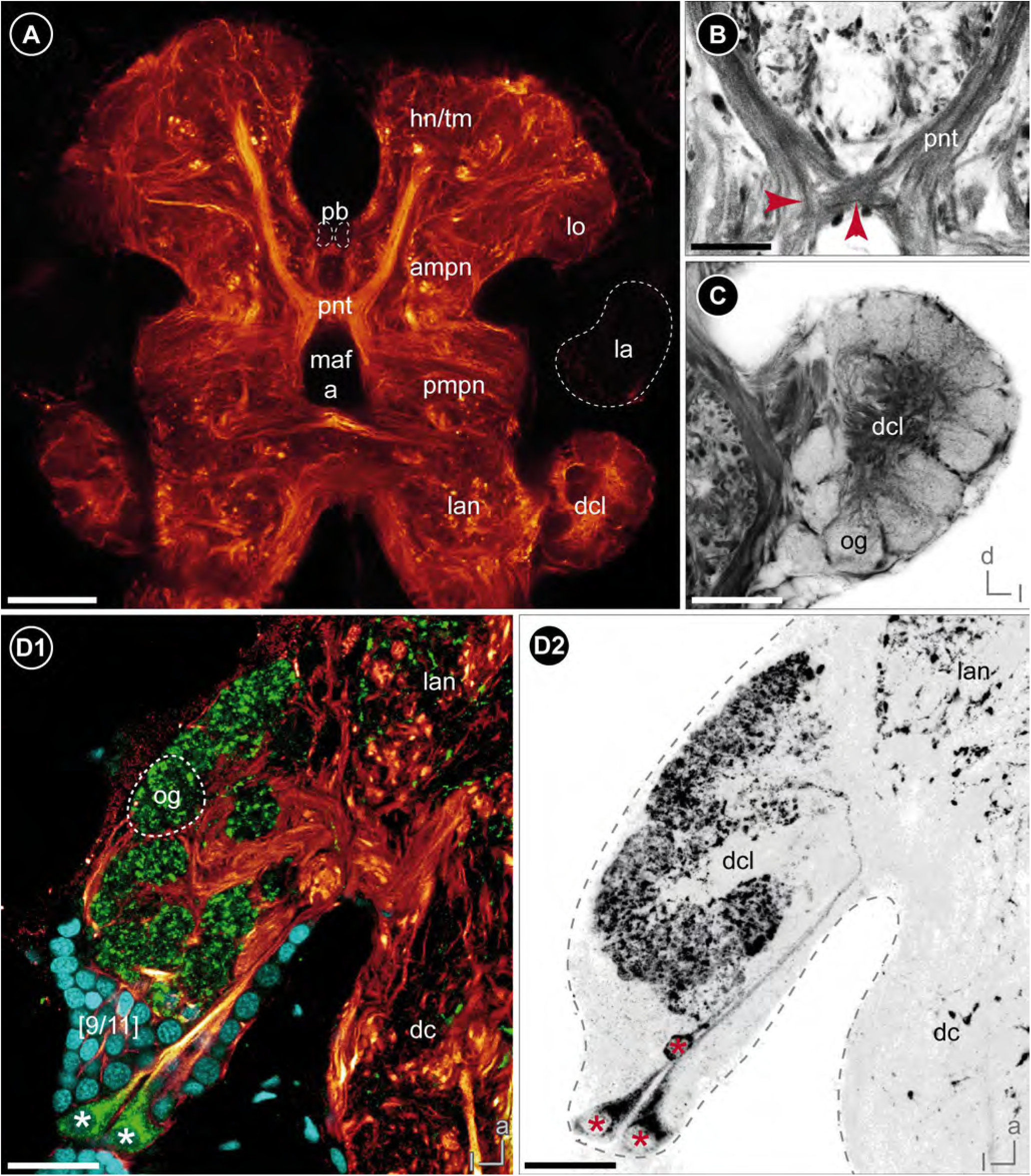
The deutocerebrum and central olfactory system. (**A**) Frontal vibratome section of the posterior part of the brain showing the projection neuron tract, labeled against acetylated tubulin. (**B**) Magnification of the chiasm of the projection neuron tract with neurites projecting contra- and ipsilaterally (red arrows), labeled against acetylated tubulin (black-white inverted). (**C**) Magnification of the deutocerebral chemosensory lobe of the right brain hemisphere showing the peripheral arrangement of spherical to wedge-shaped glomeruli, labeled against acetylated tubulin (black-white inverted). (**D**) Horizontal view on the deutocerebral chemosensory lobe of the right brain hemisphere showing three RFamide-like immunoreactive neurons located in cluster 9/11 (asterisks), triple-labeled against acetylated tubulin (yellow-red), RFamide (green) and nuclei (cyan, D1) as well as against RFamide, separately (black-white inverted, D2). *<u>Abbreviations:</u> dc* deutocerebrum, *dcl* deutocerebral chemosensory lobe, *hn/tm* hemiellipsoid body/ terminal medulla complex, *la* lamina, *lan* lateral antenna 1 neuropil, *lo* lobula, *maf a* myoarterial formation a, *og* olfactory glomeruli, *pb* protocerebral bridge, *pmpn* posterior medial protocerebral neuropil, *pnt* projection neuron tract. *<u>Scale bars:</u>* (A) 50 µm, (B-D) 25 µm.

**Fig. 6:**
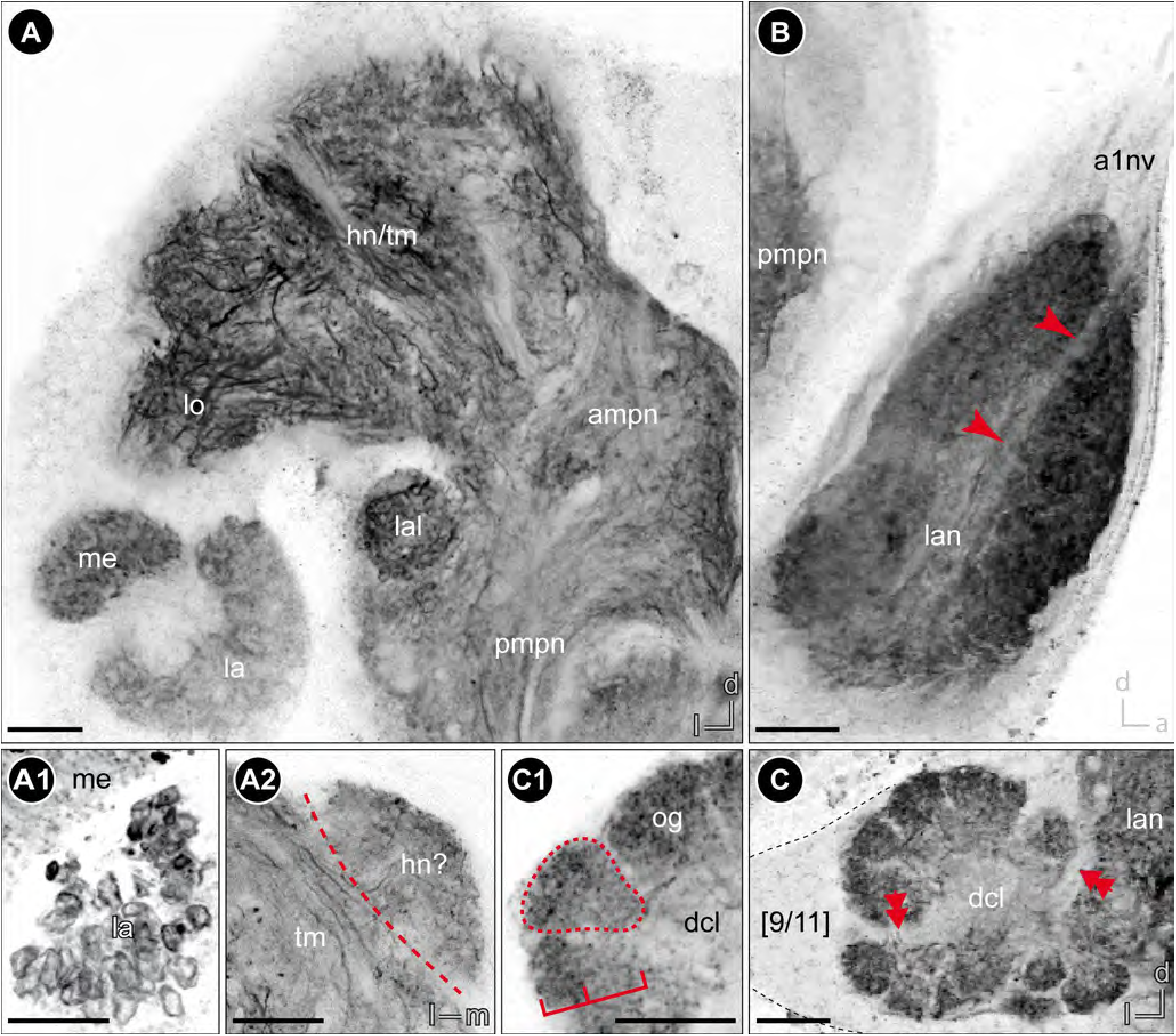
Synapsin labeling of the brain of *Parhyale hawaiensis*. (**A**) Frontal vibratome section of the anterior part of the lateral and medial protocerebrum of the brain hemisphere. (A1) Magnification of the lamina showing the organization in spherical subunits with peripherally concentrated immunoreactivity. (A2) Cross section through the dorsomedial part of the lateral protocerebrum showing a weak distinction of a cap-like structure that potentially represent the hemiellipsoid body. (**B**) Sagittal vibratome section of the lateral antenna 1 neuropil showing a weak horizontal division into two equally sized parts. (**C**) Frontal vibratome section of the deutocerebral chemosensory lobe of the left brain hemisphere. The medial and lateral foramina are discernable (double-arrows). (C1) Magnification of the olfactory glomeruli showing the spherical to wedge-shaped appearance as well as a weak division into cap and base (brackets). *<u>Abbreviations:</u> ampn* anteromedial protocerebral neuropil, *dcl* deutocerebral chemosensory lobe, *hn/tm* hemiellipsoid body/ terminal medulla complex, *la* lamina, *lal* lateral accessory lobe, *lan* lateral antenna 1 neuropil, *lo* lobula, *me* medulla, *og* olfactory glomeruli, *pmpn* posteromedial protocerebral neuropil, *[numbers]* somata cluster. *<u>Scale bars:</u>* 25 µm.

#### Median protocerebrum

The median protocerebrum is composed of the anteromedial protocerebral neuropil (ampn) and, ventrally adjacent, the posteromedial protocerebral neuropil (pmpn) that is medially divided by the myoarterial formation A (Figs. 3A, 4A). In contrast, the anteromedial protocerebral neuropil is medially fused and encloses the central complex, composed of protocerebral bridge (pb), the unpaired central body (cb), and paired lateral accessory lobes (lal). Dorsomedially, the protocerebral bridge (Fig. 5A) exhibits synapsin-like immunoreactivity and consists of two medially divided subunits that extend from anteromedial to posterolateral in a bow-like fashion (data not shown, Fig. 8). The spindle-shaped central body is located at the interface of anteromedial and posteromedial protocerebral neuropil (Figs. 1B, 3A, 4A; Add. files 2 and 3) and consists of nine horizontally arranged columnar-like subunits. These subunits display a strong RFamide-like immunoreactivity mainly in their dorsal regions. Hence, the central body appears to be divided transversally into a thinner dorsal (upper) unit and a thicker ventral (lower) unit (Figs. 3B1, 4A, 4E1). Four pairs of small tracts (w, x, y, and z) proceed from dorsal to the central body, with crossing neurites of the medial z tracts within the central body (Fig. 4E). Laterally to the central body, the lateral accessory lobes (Fig. 6A) display a weak RFamide-like immunoreactivity. They are connected by a thick protocerebral commissure in which the central body is embedded (Figs. 3A, 4A).

**Fig. 7:**
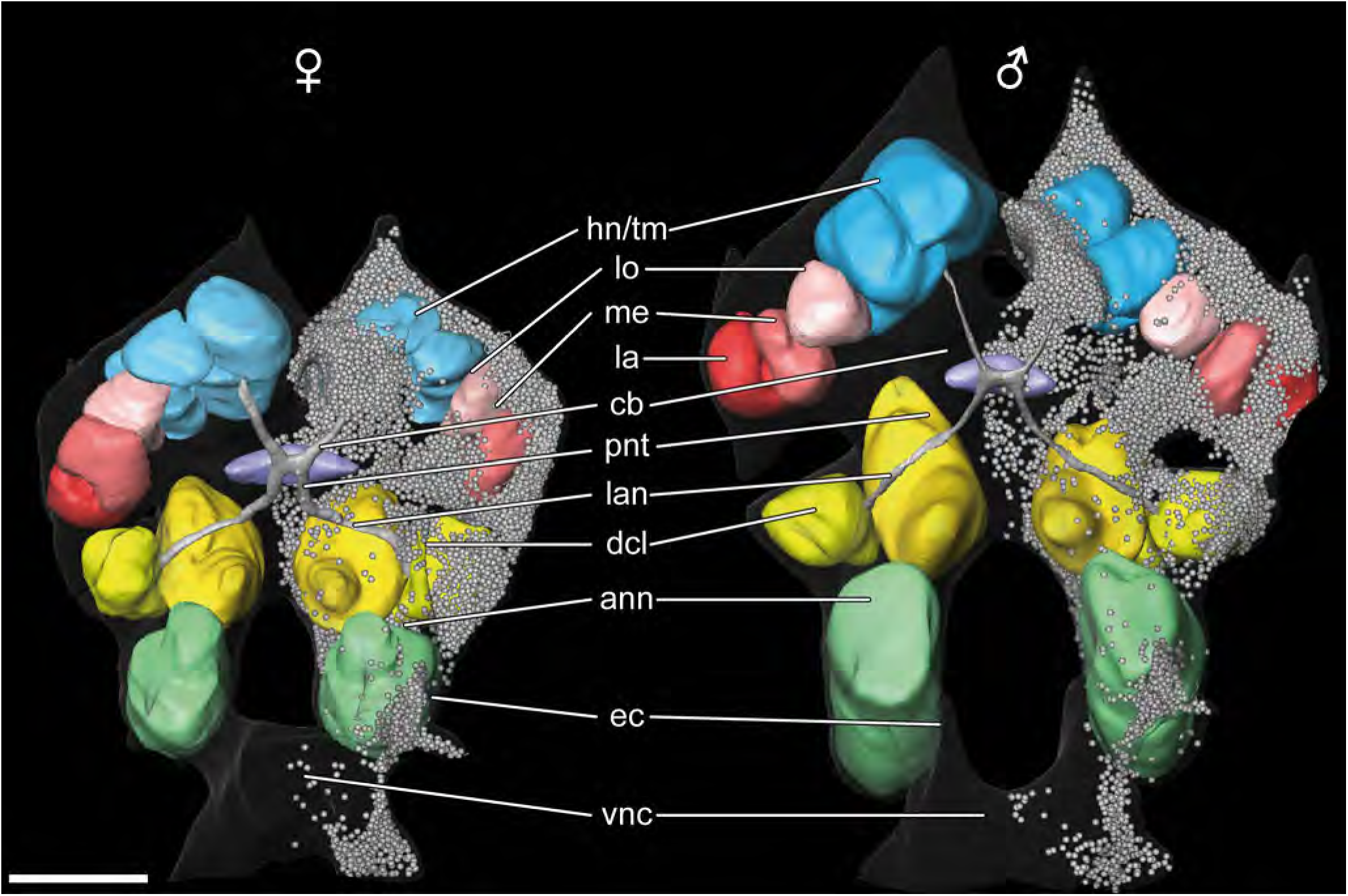
Three-dimensional reconstructions of the brains of one female and one male based on histological sections. Neuropils of the protocerebrum (reddish: visual neuropils, blue: hemiellipsoid body/terminal medulla complex, purple: central body), deutocerebrum (yellowish) and tritocerebrum (green) as well as the projection neuron tract (grey) are shown. In the right hemispheres of both brains, all somata are depicted. Interactive 3D PDFs are available as supplements (see Additional files 2 and 3). *<u>Abbreviations:</u> ann* antenna 2 neuropil, *cb* central body, *dcl* deutocerebral chemosensory lobe, *ec* esophageal connective, *hn/tm* hemiellipsoid body/terminal medulla complex, *la* lamina, *lan* lateral antenna 1 neuropil, *lo* lobula, *me* medulla, *pnt* projection neuron tract, *vnc* ventral nerve cord. *<u>Scal bars:</u>* 100 µm.

**Fig. 8:**
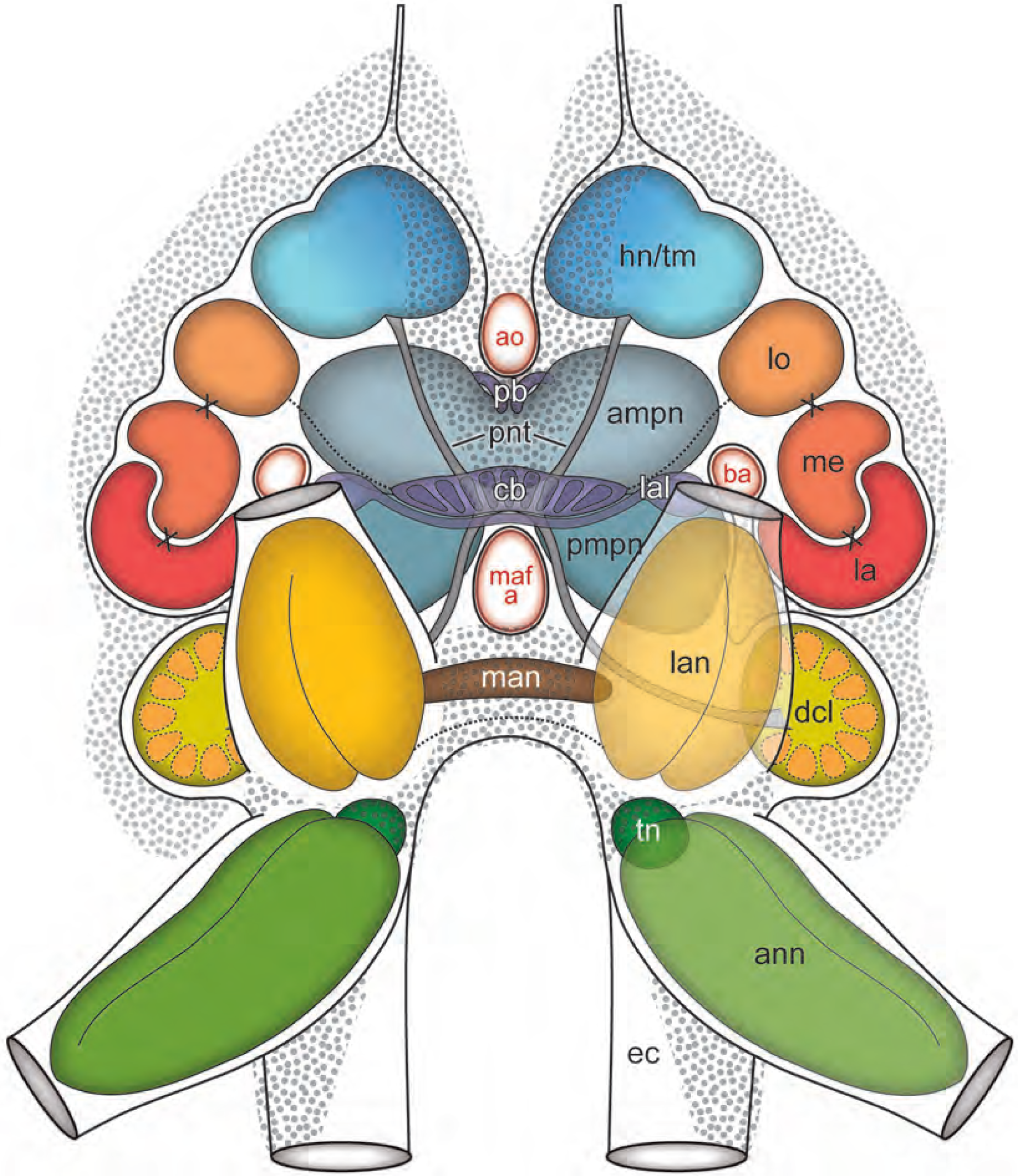
Schematic representation of the brain of *Parhyale hawaiensis* in anterior view. *<u>Abbreviations:</u> ampn* anterior medial protocerebral neuropil, *ann* antenna 2 neuropil, *ao* anterior aorta, *ba* brain artery, *cb* central body, *dcl* deutocerebral chemosensory lobe, *ec* esophageal connective, *hn/tm* hemiellipsoid body/ terminal medulla complex, *la* lamina, *lal* lateral accessory lobe, *lan* lateral antenna 1 neuropil, *lo* lobula, *maf a* myoarterial formation a, *man* medial antenna 1 neuropil, *me* medulla, *pb* protocerebral bridge, *pmpn* posterior medial protocerebral neuropil, *pnt* projection neuron tract, *tn* tegumentary neuropil.

### Deutocerebrum

The deutocerebrum is horseshoe-shaped and composed of a bipartite anterior part and a medially fused posterior part, located ventrally to the posteromedial protocerebrum. The anterior part is characterized by the large lateral antenna 1 neuropil (lan, Figs. 1C, 3, 6B) and the deutocerebral chemosensory lobe (dcl, also termed ‘olfactory lobe’, Figs. 1B, C, 2D, 3A, 5C, D, 6C; Add. files 2 and 3). In anti-SYNORF 1 synapsin labelings as well as in histological sections, the lateral antenna 1 neuropil displays a horizontal division into two equally sized parts (Fig. 6B). Posterolaterally attached to the lateral antenna 1 neuropil, the deutocerebral chemosensory lobe is innervated by a small lateral section of the antenna 1 nerve (arrows in Fig. 2D). Each deutocerebral chemosensory lobe is composed of approximate 20–40 subunits, the olfactory glomeruli (og; Figs. 3A, 5C, D, 6C). In histological sections, individual glomeruli are only weakly delineated (Fig. 2D). However, synapsin- and tubulin-like immunoreactivity reveal a multi-lobed architecture of the deutocerebral chemosensory lobe with spherical to wedge-shaped olfactory glomeruli radially arranged around a core-region that is composed of neurites of local interneurons and projection neurons (Figs. 5C, 6C). Unfortunately, a precise determination of the number of olfactory glomeruli is not possible with the used methods due to the lack of clear delimitations. In anti-synapsin labelings, the olfactory glomeruli are weakly divided into cap- and base-region (brackets in Fig. 6C1), and they display strong RFamide-like immunoreactivity across their entire volume (Fig. 5D). In our preparations, we found three characteristic local interneurons posterior to the deutocerebral chemosensory lobe, featuring strong RFamide-like immunoreactivity (asterisks in Fig. 5D). Their neurites project from the posteromedial edge of cluster 9/11 into the core-region and innervate an unknown number of glomeruli (Fig. 5D2). Neurites of laterally located somata enter the deutocerebral chemosensory lobe through a lateral foramen (double-arrow in Fig. 6C), whereas neurites of ventrally located somata bundle up medially and proceed dorsally towards the medial foramen (data not shown). Neurites of projection neurons leave the core-region through the medial foramen (double-arrow in Figs. 2D, 6C) and form a strong projection neuron tract (pnt; also termed ‘olfactory globular tract’) that proceeds dorsally to the hemiellipsoid body/ terminal medulla complex (Figs. 3A, B, 4A, 5A, 7, 8; Add. files 2 and 3). Posteriorly to the central body, the projection neuron tract forms a chiasm with ipsi- and contralateral proceeding neurites (Fig. 5A, arrows in Fig. 5B). The posteromedian deutocerebrum exhibits diffuse RFamide-like immunoreactivity (Fig. 3A). In anti-SYNORF 1 synapsin labelings, the cigar-shaped median antenna 1 neuropil (man) is located dorsally in the posterior deutocerebrum (data not shown, Fig. 8). Ventrally to the median antenna 1 neuropil and between the lateral antenna 1 neuropils, a strong deutocerebral commissure is discernible (Fig. 1B, 8).

### Tritocerebrum

The tritocerebrum is bipartite and located in close association to the circumesophageal connectives (Figs. 1B, C, 2D, 3A). It is dominated by the distinct, anteriorly oriented antenna 2 neuropil (ann, Fig. 3B, Add. files 2 and 3). In histological sections and immunohistochemical labelings, a weak horizontal division into two equally sized parts is discernable (data not shown). The innervating antenna 2 nerve (a2nv) branches off into several bundles of neurites that proceed through both halves of the antenna 2 neuropil (Fig. 3A and B). Posteromedially to the upper half of the antenna 2 neuropil, synapsin- and tubulin-like immunoreactivity reveal a small spherical neuropil, the tegumentary neuropil (tn, Fig. 3B). However, innervating tegumentary nerves could not be detected.

### Somata clusters

The neuronal somata are aggregated in clusters, which, due to their close spatial proximity, appear as a cortex that covers major parts of the brain (Fig. 1C, 7, Add. files 2 and 3). The cortex features several cell layers in different regions. As determined from two semi-thin section series, the cortex contains 6,629 nuclei (male) and 6,657 nuclei (female) per hemisphere (Fig. 7, Add. files 2 and 3). Therefore, an adult brain of *P. hawaiensis* may contain about 13,300 nuclei in total. Nonetheless, some clusters can be distinguished based on different cell body sizes and spatial relationship. The lamina is wrapped by cluster 1 that contains relatively small somata. In histological sections and immunohistochemical labelings, they appear more intensively stained than other somata (Fig. 2C, 4C1). Between hemiellipsoid body/terminal medulla complex and lobula, a distinct small cluster is located posteriorly that can be homologized with cluster 3 (Fig. 3B1). Neurites of cluster 3 proceed anteriorly into the neuropil of the lateral protocerebrum. Major parts of the median protocerebrum are covered anteriorly by cluster 6 (Fig. 3B1). Associated with the deutocerebrum, somata clusters of the central olfactory pathway are recognizable: cluster 9/11 contains somata of local interneurons and is located posterolaterally to the deutocerebral chemosensory lobe, in close proximity to cluster 1 (Fig. 5D1); somata of projection neurons form cluster 10 and are located ventrally to the deutocerebral chemosensory lobe (Fig. 2D). Between the bases of the lateral antenna 1 neuropils, the median cluster 13 is present (Fig. 2D).

## DISCUSSION

### The visual system with special emphasis on the lamina

Depending on the architecture of the compound eyes, there is a high disparity in morphology and size of visual neuropils in amphipod crustaceans [39,41]. The visual neuropils may be anteriorly oriented as in *Streetsia* sp. [59] or laterally extended as in *Platorchestia platensis* [43] and *Gammarus setosus* [41], or bent backwards as in *Orchestia cavimana* [44] and *P. hawaiensis* (this study). The sharp turn of the neuraxis within the protocerebrum of *P. hawaiensis* is not only indicated by the arrangement of visual neuropils from ventrolateral to dorsomedial, but also mirrored in the chiasm of the optical nerve. We assume that this chiasm results from the upside-down orientation of the visual neuropils and does not constitute a structural modification.

The visual neuropils comprise lamina, medulla and lobula. In amphipod brains, the lobula seems to be poorly developed and, as a consequence, is hard to distinguish from the terminal medulla [39,44], which might result from the low number of ommatidia and other neuronal elements of the visual pathway. However, there is a distinct connection between the lobula and the lateral part of the central body, that might provide visual input to the latter (reviewed in [60]). Hence, the lobula can be identified as a spherical neuropil in close association with both, terminal medulla and medulla. The outer chiasm (between lamina and medulla) is present in all previously investigated amphipod species as well as in *P. hawaiensis* [33,38,41,44]. In contrast, the presence of an inner chiasm (between medulla and lobula) was mostly denied in amphipods [33,61], which might be due to the close proximity of medulla and lobula and an uncertain delineation of the latter. This suggestion is support by investigations of brains of oxycephalid, hyperiid, phronimid, auchylomerid and tryphenid amphipods [59], which possess well-differentiated visual neuropils (lamina, medulla, and lobula) as well as outer and inner chiasmata. In sagittal sections, we detected a few neurites crossing each other in the corresponding region between medulla and lobula, an arrangement that matches the characteristics of the malacostracan inner chiasm (compare [62,63]). Thus, we suggest the presence of an inner chiasm and lobula in agreement with the suggested ground pattern of the malacostracan brain [57], which contradicts Ramos et al. [33], who stated that the inner chiasm is absent and the third visual neuropil represents the lobula plate.

In *P. hawaiensis*, medulla and lobula appear as homogenous structures whereas the lamina shows specific structural features. It consists of numerous spherical subunits that display synapsin-like immunoreactivity only peripherally and, in anti-histamine labelings, shows a two-layered organization. In most previous studies on amphipod brains, the specific structure of visual neuropils was not described. Only Gräber [38] recognized a ‘distinctly granular’ (“deutlich gekörnelt”) structure of the lamina in *Gammarus* spp. Tyrosinated tubulin-like immunoreactivity revealed transversely projecting neurites within the lamina of *O. cavimana* ([44], their Fig. 2B), arranged in a pattern that corresponds to the typical, retinotopic organization of laminae in crustaceans [64]. However, the chambered appearance of the *P. hawaiensis* lamina has not been described in any amphipod or other malacostracan crustaceans so far, and does not represent the typical organization of optic cartridges. Using opsin transgene reporters to visualize photoreceptor projections, Ramos et al. [33] revealed similar profiles in the lamina of *P. hawaiensis*. Thus, we suggest that this chambered appearance results from specialized photoreceptor terminals. Comparable to *P. hawaiensis*, Sombke and Harzsch [65] identified club-shaped terminations of histamine-like immunoreactive photoreceptor cell axons in the lamina of the centipede *Scutigera coleoptrata* that exhibit a chambered appearance, which is however not present in anti-synapsin labelings.

### Do amphipod brains possess a hemiellipsoid body?

In the malacostracan brain, the (bilaterally paired) hemiellipsoid body and terminal medulla are targeted by axons of the olfactory projection neurons as output pathway of the olfactory lobe and suggested to function as higher integrative centers (among others) (reviews [36,37,66]). In all previous studies on amphipod brains, the authors mentioned and discussed the absence of the hemiellipsoid body [38,39,41,43,44,67]. Furthermore, Ramm and Scholtz [44] questioned the suitability of previously used characteristics to identify the hemiellipsoid body in Peracarida, including in the two isopod species *Saduria entomon* and *Idotea emarginata*, in which the presence of the hemiellipsoid body had previously been assumed [57,68]. In this context, Ramm and Scholtz [44] promoted the suggestion by Stegner et al. [67] that the loss of the hemiellipsoid body might be an apomorphic feature of a subgroup of Peracarida (Amphipoda + Mancoida *sensu lato*) and the terminal medulla may function as the only second order olfactory center. However, the hemiellipsoid body that was previously described as part of the terminal medulla (as region I and II, [69]) is known to show various grades of complexity in different species. In decapod crustaceans, the structural complexity of the hemiellipsoid body, e.g. its regionalization into cap and core, coincides with the number of antennular afferents, the subdivision of olfactory glomeruli, and the number of inter- and projection neurons [37,70,71]. Correspondingly, decapod crustaceans with an elaborate olfactory system display a highly pronounced hemiellipsoid body (e.g. in *Birgus latro*, [71]. *Vice versa*, in dendrobranchiate Decapoda, e.g. *Penaeus duorarum* and *P. vannamei,* in which a subdivision of olfactory glomeruli is only indicated by immunohistochemical experiments, the lateral protocerebrum is poorly differentiated and the hemiellipsoid body hard to detect [72] and sometimes only described in a complex with the terminal medulla [73]. Another critical point is that the closely associated terminal medulla is a highly complex and not precisely defined neuropil region that cannot be delimited from other neuropils with certainty. Therefore, defining the expected morphological structure of the hemiellipsoid body is a crucial step to discuss its presence or absence.

In *P. hawaiensis*, the olfactory glomeruli are only weakly differentiated. Because of this observation together with the relatively low number of glomeruli and olfactory interneurons when compared to other malacostracans [37], we expect *a priori* a poorly differentiated hemiellipsoid body. In the dorsomedial region of the lateral protocerebrum, synapsin-like immunoreactivity reveals a cap-like structure that, in its degree of differentiation, may correspond to the hemiellipsoid body of e.g. *Penaeus duorarum* [72]. However, tubulin-like and RFamide-like immunoreactivity indicate a division of a larger region than this cap-like structure in *P. hawaiensis*. Therefore, we are not able to define a hemiellipsoid body with certainty. Nevertheless, as the hemiellipsoid body is part of the ground pattern of the malacostracan brain [57] and a subdivision is indicated by immunohistochemical labelings, we suggest its presence in the brain of *P. hawaiensis.* Furthermore, the hemiellipsoid body may be generally accepted, at least in complex with the terminal medulla, in amphipod brains. Considering that in malacostracan crustaceans the hemiellipsoid body seems to function as a multimodal integrative center and has been suggested to function in learning and memory [62,74,75], we conclude that such functions play only subordinate roles in the behavioral repertoire of amphipods.

### Amphipods and the organ of Bellonci

In *P. hawaiensis*, the hemiellipsoid body/terminal medulla complex is innervated by a paired small nerve from dorsal. In the amphipod genera *Caprella* and *Gammarus*, Hanström [76] and Gräber [38] likewise described a pair of protocerebral nerves that fuse to a single nerve (in *Gammarus* spp.) and connect the frontal organ with the terminal medulla. However, the term ‘frontal organ’ was not clearly defined in the past and used for different structures by different authors. Therefore, the term ‘organ of Bellonci’ was reintroduced to differentiate between photoreceptive organs and “other types of receptors innervated from the medullae terminales” [77]. Although the identification of the organ of Bellonci was related to some morphological criteria (extracellular cavity and ciliated neurons, [77–79]), terminological inaccuracies remained. For example, the originally described frontal organ of Branchiopoda, which was also termed ‘X-organ’ or ‘organ of Bellonci’, was reinterpreted as ‘frontal filament organ’ [80]. In contrast to the organ of Bellonci, the frontal filament organ is composed of bipolar neurons connecting the frontal filament and the lateral protocerebrum.

Based on microCT analysis, we detected the organ of Bellonci that is connected to the median protocerebrum via a paired small nerve, and which in shape and position is similar to isopods and other amphipods [81–83]. In malacostracan crustaceans, the organ of Bellonci displays considerable morphological variations [84]. It commonly consists of one or two sensory pores and afferent neurites that innervate the so-called ‘onion bodies’ that are in close proximity to the visual neuropils and the hemiellipsoid bodies (e.g. [84,85]). Ultrastructural studies suggested a mainly sensory function [79,84,86], but the sensory modality is still a matter of debate [84,85].

### The central complex

All major arthropod taxa possess a protocerebral midline neuropil called central body that in hexapods and crustaceans is part of the central complex [56,87]. Several lines of evidence from behavioral and comparative anatomical and physiological studies suggest that the central complex serves as a motor control center that is involved in orchestrating limb actions. The composition of protocerebral bridge, central body, paired lateral accessory lobe and four pairs of tracts (w, x, y, and z) present in *P. hawaiensis* resembles the central complex as described in crayfish and many other Malacostraca [35,87,88]. In *P. hawaiensis*, the protocerebral bridge consists of two arched neuropils that are medially separated and arranged in a V-shape. Except for the tripartite protocerebral bridge in *Gammarus* spp. (including one median, unpaired element [38]), the protocerebral bridge of amphipod crustaceans generally consists of two medially divided halves that are connected by a small commissure (summarized in [39]). In the amphipod species *O. cavimana* and *N. puteanus*, this commissure was shown in serotonin-like immunoreactive labelings [44], which was not performed in the present study. Anteroventrally to the protocerebral bridge, the central body receives input from cluster 6 by four pairs of tracts, termed w, x, y, and z. These tracts are formed by columnar neurites that cross the midline only within the central body, what is suggested to be part of the malacostracan ground pattern [35]. While the central complex of *P. hawaiensis* exhibits all four pairs of tracts, there were only three pairs identified in *N. puteanus* and none in *O. cavimana* [44]. Besides different numbers of interconnecting tracts, the number of horizontally arranged subunits of the central body also differs in amphipod crustaceans. Similar to *P. hawaiensis*, the genus *Vibilia* features nine horizontally arranged subunits [39], whereas seven subunits were described in *Gammarus* spp. [38,41] and only four in *N. puteanus* [44]. In contrast, species of the genera *Orchestia* and *Phronima* display homogenous central bodies without subdivisions [39,43,44]. In *P. hawaiensis, N. puteanus,* and *O. cavimana*, the number of subunits of the central body decreases with a decreasing number of interconnecting tracts. The horizontal subdivision of the central body may be dependent on the number of tracts, as previously suggested by Homberg [60], although the numbers are not equal. Anyway, Ramm and Scholtz [44] as well as Stegner et al. [67] discussed a possible correlation between lack of eyes and a decreasing number of tracts that seems to comply with our findings in *P. hawaiensis*.

### The central olfactory pathway

The deutocerebral chemosensory lobes receive input from unimodal chemosensory sensilla on the first antennae, the aesthetascs (reviewed in [37]). The afferents of chemosensory receptor neurons associated with bimodal sensilla are assumed to mainly innervate the lateral antenna 1 neuropil, if not exclusively [89]. Nonetheless, in *P. hawaiensis, Caprella acutifrons*, and *Gammarus* spp., afferents of olfactory sensory neurons associated with the aesthetascs form a distinct bundle in the lateral part of the nerve of the first antenna [38,90]. These afferents innervate the olfactory glomeruli, which are spherical in *N. puteanus* [44] and slightly more wedge-shaped in *Gammarus* spp. [38] and *P. hawaiensis*. Spherical olfactory glomeruli are part of the ground pattern of malacostracan brains and display a plesiomorphic feature [56,57,91,92]. For decapod crustaceans, it was already suggested that an increasingly elongated shape of olfactory glomeruli coincides with an increasing complexity of the olfactory system [70,71,73]. In the brain of *P. hawaiensis*, anti-synapsin labelings reveal a slight regionalization of the olfactory glomeruli into cap and base that is known from many malacostracans where olfactory glomeruli can be differentiated in up to three regions (cap, subcap, and base). This regionalization is a consequence of region-specific interconnection of afferents and different types of interneurons [37,93]. Hence, the cap represents the major input region where afferents terminate, whereas the base represents the major output region where dendrites of projection neurons arborize. Neurites of local interneurons terminate in both regions. In *P. hawaiensis*, somata of local interneurons (cluster 9/11) encompass posterolaterally the deutocerebral chemosensory lobe and house three RFamide-like immunoreactive neurons. RFamides are known to characterize local interneurons in the olfactory glomeruli of numerous decapods, e.g. *Panulirus argus* [94] and *Penaeus vannamei* [73]. Another cluster of somata is located ventrally to the deutocerebral chemosensory lobes and, in accordance with *N. puteanus* [44], is suggested to represent cluster 10, which houses the projection neurons. Their neurites form a strong tract that projects dorsally towards the medial foramen of the deutocerebral chemosensory lobe. We assume that both, dendrites entering the deutocerebral chemosensory lobe and axons leaving it, pass through the medial foramen. In comparison to other malacostracans, the deutocerebral chemosensory lobe with its low number of olfactory glomeruli displays a little complex architecture [37]. Thus, it is conceivable that the olfactory landscape, so the number of detectable substances, might be rather small in *P. hawaiensis*.

### Other deutocerebral neuropils and tritocerebrum

Besides the deutocerebral chemosensory lobe, the median and lateral antenna 1 neuropils are principal components of the deutocerebrum in Malacostraca [34,57,87]. The median antenna 1 neuropil processes primarily input from proximal antennal segments, in particular from the statocyst [95,96], and were also described in *O. cavimana* and *N. puteanus* [44]. The lateral antenna 1 neuropil receives mechanosensory input from bimodal sensilla on the first antenna [36,37,97] and contains dendrites of motor neurons controlling the movement of the first antenna [98,99]. In *P. hawaiensis*, the lateral antenna 1 neuropil displays a bi-lobed shape as in the stomatopod *Neogonodactylus oerstedii* [100] and in many decapods [54] that was also described in *N. puteanus*, but not in *O. cavimana* [44]. The tritocerebrum of *P. hawaiensis* is dominated by the antenna 2 neuropil, which receives mechanosensory input from the second antenna and controls its movements [101,102]. Both neuropils, the lateral antenna 1 neuropil and the antenna 2 neuropil, are known to show a stratified appearance corresponding to a somatotopic organization in many malacostracans [73,100,103]. However, a stratification is not part of to the ground pattern of malacostracans [57] and not evident in amphipod brains ([44], this study). A tritocerebral tegumentary neuropil is present medially to the antenna 2 neuropil in *P. hawaiensis, G. setosus* [41], and *O. cavimana*, but seems to be absent in *N. puteanus* [44]. The tritocerebral tegumentary neuropil receives input from the paired tegumentary nerve that transmits mechanosensory information from the carapace (not from the organ of Bellonci). Although this nerve could not be identified with certainty, it was previously described by Divakaran [45] in *P. hawaiensis*.

### Sexual dimorphism in amphipod brains?

Amphipod crustaceans show sexual dimorphisms [104]. Matured amphipod males are often larger than females, but the most striking feature is the significantly enlarged second pereopod in males (i.e. gnathopod; Fig. 1A) [104,105]. In laboratory conditions, it is noticeable that males are more actively moving around by leaving the coral gravel more frequently (personal observation). They also constantly check nearby females about their fertility status to grab receptive females in precopula (Fig. 1A) for later sexual reproduction. Investigating the reproductive behavior of the amphipod species *Gammarus palustris*, Borowsky and Borowsky [106] assumed that contact pheromones might play an important role for this behavior. These speculations fit to descriptions on sexual dimorphisms of the peripheral sensory system that first antennae of male amphipods can be equipped with male-specific sensilla [107,108]. However, the presence and sensory processing of possible attractants is still unresolved. Sexual dimorphism of the deutocerebral chemosensory lobe was described in Mysidacea and Euphausiacea [109] and similarly suggested in Brachyura [110]. Slight differences were also noticed in the central bodies of male and female individuals of the brachyuran genus *Uca* [111]. However, based on our data set we used we were not able to detect sexual dimorphism in the brain of *P. hawaiensis*. Therefore, further investigations are required to discover possible neuroanatomical foundation for sexual dimorphic behavior in *P. hawaiensis*.

## CONCLUSIONS

In comparison to the ground pattern of Malacostraca [57], the brain of *Parhyale hawaiensis* exhibits all neuropils of the proto-, deuto-, and tritocerebrum with exception of lobula plate (protocerebrum) and projection neuron tract neuropil (deutocerebrum). Some neuropils were difficult to demarcate due to a close association to each other and uniform appearance (such as lobula, terminal medulla, and hemiellipsoid body), which seems to be a common feature of Amphipoda [38,43,44]. Beyond the uniformity of amphipod brains, there is also a certain degree of variability in architecture and size of different neuropils [59]. Ramm and Scholtz [44] suggested that size and structural elaboration of neuropils might correlate with ecology and life style in different species. For example, the cave-dwelling amphipod species *N. puteanus* possesses reduced visual neuropils corresponding to its reduced compound eyes, and the terrestrial amphipod species *O. cavimana* possesses reduced deutocerebral chemosensory lobes in correspondence to reduced first antennae (compare also [43]). Amphipod species from deep-sea feature either well-developed sense organs and associated neuropils of the chemosensory or visual system [59]. The brain of *Streetsia* sp. is dominated by well-differentiated visual neuropils in correspondence to its large eyes and the brain of *Corophium volutator*, whose second antennae are considerably enlarged, is characterized by a pronounced tritocerebrum [59]. In summary, we conclude that the obvious disparity of amphipod life histories and ecologies is clearly mirrored in their neuroanatomy. As the brain of *P. hawaiensis* does not display striking modifications from the suggested ground-pattern of Malacostraca or any bias towards one particular sensory modality, we assume that its brain may represent the common type of the amphipod brain.

## ABBREVIATIONS

a: anterior
a1: antenna 1
a1nv: antenna 1 nerve
a2: antenna 2
a2nv: antenna 2 nerve
ampn: anterior medial protocerebral neuropil
ann: antenna 2 neuropil
ao: anterior aorta
ba: brain artery
br: brain
cb: central body
dc: deutocerebrum
d: dorsal
dcc: deutocerebral commissure
dcl: deutocerebral chemosensory lobe
ec: esophageal connectives
es: esophagus
g: segmental ganglion (**1-7** of the pereon, **I-VI** of the pleon)
gn: gnathopod
hn/tm: hemiellipsoid body/terminal medulla complex
l: lateral
la: lamina
lal: lateral accessory lobe
lan: lateral antenna 1 neuropil
lo: lobula
maf a: myoarterial formation a
man: medial antenna 1 neuropil
me: medulla
og: olfactory glomeruli
om: ommatidia
onv: optical nerve
p: pereopod
pb: protocerebral bridge
pc: protocerebrum
pcc: protocerebral commissure
pl: pleopod
pmpn: posterior medial protocerebral neuropil
pnt: projection neuron tract
re: retina
seg: subesophageal ganglion
tc: tritocerebrum
tn: tegumentary neuropil
vnc: ventral nerve cord

## DECLARATIONS

### Acknowledgements

The authors thank Jakob Krieger, Tina Kirchhoff, Peter Wagenknecht, Elisabeth Böttcher, Marie K. Hörnig (University of Greifswald) and Joris Wiethase (Humboldt University of Berlin) for technical support. We are also grateful to Nicholas J. Strausfeld (University of Arizona, Tucson) and several other discussion partners at conferences and workshops for helpful exchange of thoughts.

### Availability of data and materials

The data generated and/or analyzed during the current study are available from the corresponding authors upon reasonable request.

### Authors’ contribution

All authors had full access to all data in the study and take responsibility for the integrity of data and the accuracy of data analysis. Study concept and design: ChW, CaW, AS. Acquisition of data: ChW, AS. Analysis and interpretation of data: ChW, CaW, SH, AS. ChW drafted the manuscript with further input from all co-authors.

### Ethics approval and consent to participate

Ethical approval and consent to participate were not required for this work.

### Consent for publication

Not applicable.

## FIGURES AND ADDITIONAL FILES

**Additional file 1:**
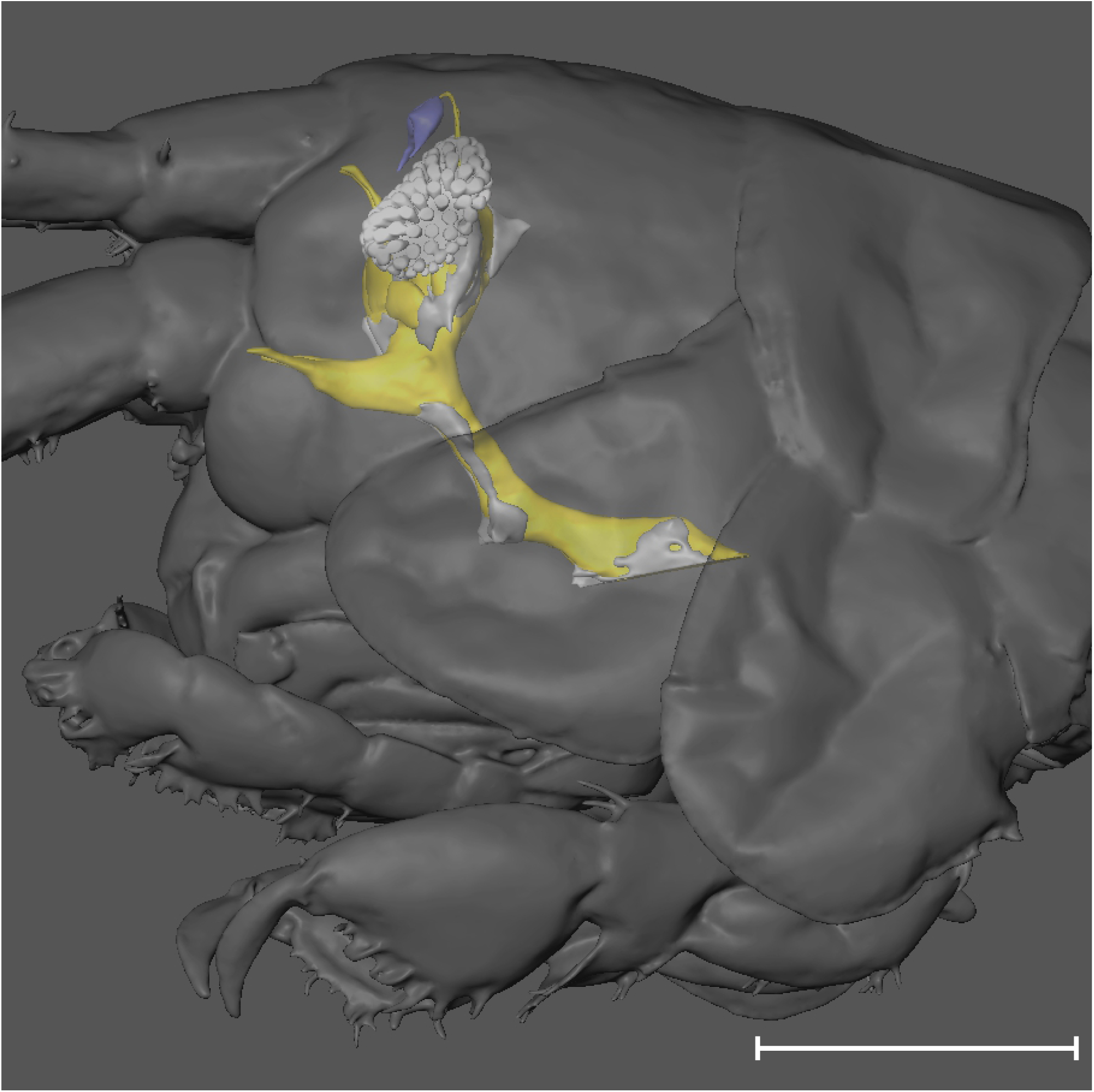
Interactive content related to figure 1C.Three-dimensional reconstruction of a male brain located within the head capsule of *P. hawaiensis* based on microCT data. The PDF version contains interactive 3D content. To activate, click on the figure in Adobe Reader and by using the computer mouse you can bring the model in any desired position and magnification. Using the model hierarchy, you can in-or exclude all different brain components. For further functionalities see the content menu. Color code: yellow: brain neuropil, gray: brain cortex, white: optic system, blue: organ of Bellonci. Scale bar: 500µm

**Additional file 2:**
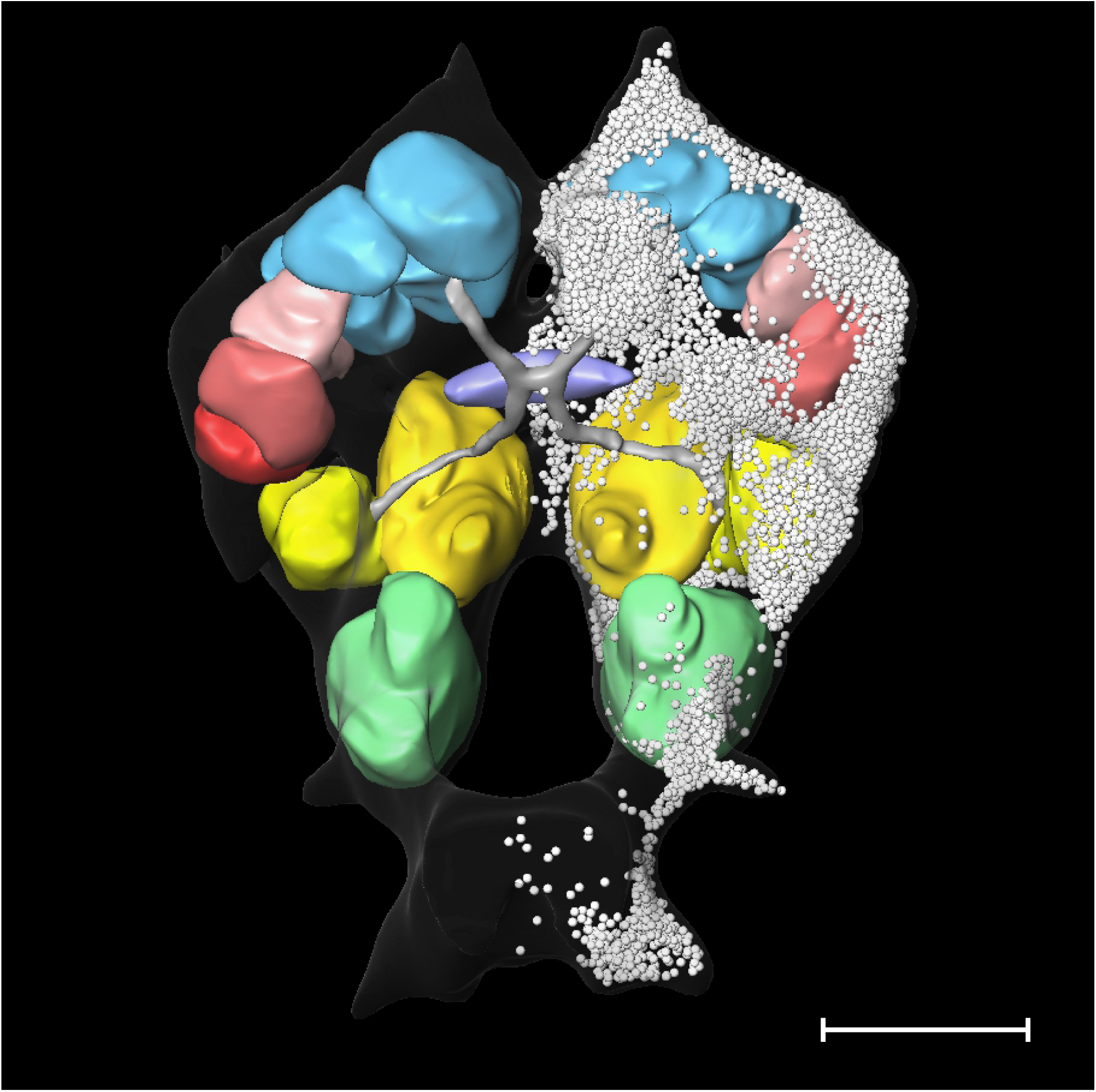
Interactive content related to figure 7. Three-dimensional reconstruction of a female brain of *P. hawaiensis* based on histological sections. The PDF version contains interactive 3D content. To activate, click on the figure in Adobe Reader and by using the computer mouse you can bring the model in any desired position and magnification. Using the model hierarchy, you can in-or exclude all different brain components. For further functionalities see the content menu. Abbreviations: ann antenna 2 neuropil, cb central body, dcl deutocerebral chemosensory lobe, hn/tm hemiellipsoid body/ terminal medulla complex, la lamina, lan lateral antenna 1 neuropil, lo lobula, me medulla, pnt projection neuron tract. The nuclei of the brain (somata) only shown in the right hemisphere. Scale bar: 100µm

**Additional file 3:**
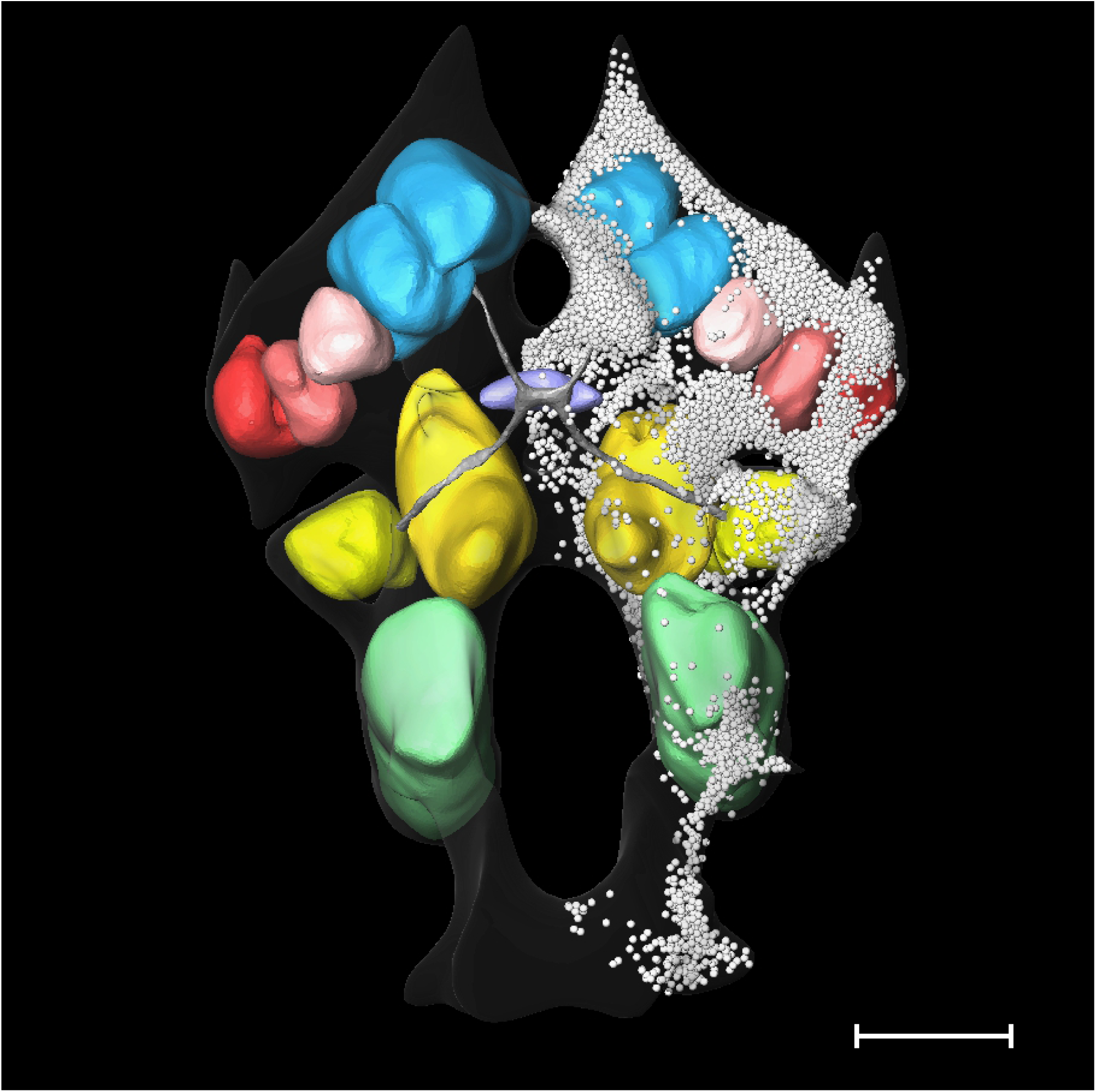
Interactive content related to figure 7. Three-dimensional reconstruction of a male brain of *P. hawaiensis* based on histological sections. The PDF version contains interactive 3D content. To activate, click on the figure in Adobe Reader and by using the computer mouse you can bring the model in any desired position and magnification. Using the model hierarchy, you can in-or exclude all different brain components. For further functionalities see the content menu. Abbreviations: ann antenna 2 neuropil, cb central body, dcl deutocerebral chemosensory lobe, hn/tm hemiellipsoid body/ terminal medulla complex, la lamina, lan lateral antenna 1 neuropil, lo lobula, me medulla, pnt projection neuron tract. The nuclei of the brain (somata) only shown in the right hemisphere. Scale bar: 100µm

